# Heterogeneity in the respiratory symptoms of patients with mild-moderate COPD

**DOI:** 10.1101/401695

**Authors:** Kate M. Johnson, Abdollah Safari, Wan C. Tan, Jean Bourbeau, J Mark FitzGerald, Mohsen Sadatsafavi, for the Canadian Cohort of Obstructive Lung Disease (CanCOLD) study and the Canadian Respiratory Research Network

## Abstract

**Background:** The burden of symptoms varies markedly between patients with Chronic Obstructive Pulmonary Disease (COPD) and is only weakly correlated with lung function impairment. While heterogeneity in lung function decline and exacerbations have been previously studied, the extent of heterogeneity in symptoms and the factors associated with this heterogeneity are not well understood.

**Methods:** A sample of the general Canadian population ≥40 years with persistent airflow limitation was followed for up to 3 years. Participants reported whether they experienced chronic coughing, phlegm, wheezing, or dyspnea during visits at 18-month intervals. We used mixed-effect logistic regression models (separately for each symptom) to assess overall heterogeneity in the occurrence of symptoms between individuals, and the proportion of variation in symptom burden explained by lung function versus all other clinical characteristics of participants.

**Results:** 548 participants (54% male, mean age 67 years) contributed 1,086 visits in total, and 82% of patients reported at least one symptom during follow-up. There was substantial heterogeneity in the individual-specific probabilities for the occurrence of symptoms. This heterogeneity was highest for dyspnea and lowest for phlegm (interquartile range of probabilities: 0.15-0.77 and <0.01-0.53, respectively). FEV_1_ explained 82% of the variation between individuals in the occurrence of phlegm, 26% for dyspnea, 3% for cough, and <0.1% for wheeze. All clinical characteristics of participants (including FEV_1_) explained between 86% of heterogeneity in the occurrence of phlegm to <1% for wheeze.

**Conclusion:** There is marked heterogeneity in the burden of respiratory symptoms between COPD patients. The ability of lung function and other commonly measured clinical characteristics to explain this heterogeneity differs between symptoms.

## 1.0 INTRODUCTION

Chronic obstructive pulmonary disease (COPD) is a common inflammatory lung condition that affects close to 400 million people worldwide.^1^ COPD is characterized by persistent airflow limitation and symptoms such as breathlessness, chronic cough, sputum production, wheezing, and chest tightness.^2^ Respiratory symptoms are a major burden in many patients, and are associated with an increased frequency of exacerbations,^3^ worse disease prognosis,^4–6^ lower health status,^7,8^ reduced quality of life,^9^ and higher healthcare resource utilization.^10^

The three major components of the natural history of COPD are lung function status, patterns of exacerbations, and symptom burden.^2^ The degree of symptom impairment is increasingly recognized as an important determinant of patient management strategies, and one that is only partially dependent on the severity of airflow limitation.^4,7,11,12^ Indeed, modern guidelines such as the Global Initiative for Chronic Obstructive Lung Disease (GOLD) appreciate the importance of all three components in disease management decisions. The GOLD guidelines recommend evaluating symptoms separately from airflow limitation and history of exacerbations in providing therapeutic recommendations.^2^

It is increasingly recognized that COPD is a heterogeneous disease. Individuals can vary markedly in their rate of lung function decline^13^ and frequency of exacerbations^14,15^ over the course of their disease. For example, COPD patients in the Lung Health Study had an annual rate of change in FEV_1_ that ranged from rapidly declining to modestly increasing (95% CI −83 mL/yr to +15 mL/yr).^13^ Similarly, the annual rate of exacerbations observed in the MACRO clinical trial varied from 0.47 to 4.22.^15^ Quantifying this variation at an individual level is critical to enabling precise risk factor and disease management.^16^

In contrast, heterogeneity in the burden of symptoms has not received the same level of attention as these other disease components. Previous studies have reported that patient symptoms tend to vary over the day, week, or season,^11,17–19^ but the extent of variation between individuals in the occurrence of symptoms has been less well characterized. Understanding the extent and drivers of this heterogeneity can help improve our understanding of the natural history of COPD and ultimately help formulate disease management strategies that provide optimal therapeutic strategies for each patient.

Using data from a population-based prospective cohort, we assessed the burden of self-reported respiratory symptoms in patients with persistent airflow limitation in order to (1) characterize variation in the occurrence of symptoms between individuals, and (2) determine the proportion of between-individual variability in symptoms that can be explained by lung function versus all other observable characteristics. We hypothesized that there is high variability in the occurrence of symptoms between individuals, and that an individual’s clinical and demographic characteristics explain a larger fraction of this heterogeneity than lung function alone.

## 2.0 METHODS

We used data from the Canadian Cohort of Obstructive Lung Disease (CanCOLD), which is a multicenter prospective longitudinal cohort study conducted across Canada.^20^ Individuals ≥40 years old were recruited using random digit dialing and multi-level sampling to ensure representativeness of the general Canadian population. Participants were followed for a maximum of 3 years with in-person visits at baseline and at 18-month intervals. From the entire cohort of CanCOLD participants (N=1,532), we selected all visits in which the patient had persistent airflow limitation, defined as post-bronchodilator FEV1/FVC< lower limit of normal (LLN).^21^

Information was collected during each visit on the presence of cough, phlegm, wheeze, and dyspnea using separate questions for each symptom. Participants reported whether they (1) usually coughed in the absence of a cold, (2) brought up phlegm from the chest in the absence of a cold, and (3) experienced any wheezing or whistling in the chest. Dyspnea (4) was measured using the Medical Research Council (MRC) dyspnea scale,^22^ which was converted to a binary variable by assuming that a score of 2-5 indicated the presence of dyspnea. Dyspnea and whether the participant experienced any symptoms were assessed in a subset of the data that included 957 visits from 502 participants because 46 participants were unable to walk and therefore did complete the MRC dyspnea test. Participants reported their demographic information, smoking status and history, number of comorbidities, and history of physician-diagnosed COPD, emphysema, or chronic bronchitis at each visit using validated questionnaires with a recall period spanning the length of time between visits.^23^

### Statistical analysis

We used separate random effect logistic regression models for cough, phlegm, wheeze, dyspnea, and any symptoms to model heterogeneity. The random effect term captured the variability among individuals (heterogeneity) that was not attributable to the independent variables in the model. We initially determined the total heterogeneity in the occurrence of symptoms using an intercept-only logistic regression model for each symptom (the null model, *ie*, no independent variables). We used this model to determine the individual-specific probability of experiencing each symptom, and estimated the interquartile (25%-75%) range of probabilities to measure heterogeneity in the occurrence of symptoms.

We subsequently assessed the proportion of the total heterogeneity in symptoms that could be explained by all measured characteristics of individuals. For this, we included patient age, sex, body mass index (BMI), ethnicity, number of comorbidities, smoking status, pack-years of smoking, post-bronchodilator forced expiratory volume in 1 second (FEV_1_), and diagnosis status as independent variables in each model (the full models). In order to determine the variance explained by the independent variables (*ie,* participants’ measured characteristics), we calculated the proportion of variance in the random effect of the full model for each symptom (with all the independent variables), compared to the variance in the random effect of the null model with no independent variables.^24^ We repeated this process using a reduced model with FEV_1_ as the only independent variable (as opposed to the full model) to determine the proportion of total heterogeneity explained by lung function alone.

We conducted sensitivity analyses in which percent predicted FEV_1_ and GOLD grade were used in place of FEV_1_ as indicators of lung function (collinearity prevented these variables from being included in the model at the same time). Seasonality was not included in the main analysis because the recall period was >1 year and therefore spanned all seasons; however, season was assessed in a sensitivity analysis to account for the possibility that patients were more likely to recall their recent symptom burden (which could be affected by the current season). All analyses were performed in SAS (version 9.4, 2016).

## 3.0 RESULTS

The characteristics of participants are shown in ***Table 1***. There were 1,086 visits from 548 participants in the final sample (54% male, mean age 67 years). 92% of participants had mild to moderate disease (grade I-II), 7% had severe disease (grade III), and 1% had very severe disease (grade IV) as measured by GOLD grades.^2^ 74% of participants with persistent airflow limitation on spirometry had not been previously diagnosed. The average follow-up time was 18 months; 39% of participants underwent only one study visit, and 37% of participants were assessed at three study visits. The characteristics of the subset of the data used to analyze dyspnea and any symptoms were very similar (***Supplementary Material, Table 1***).

**Table 1.**
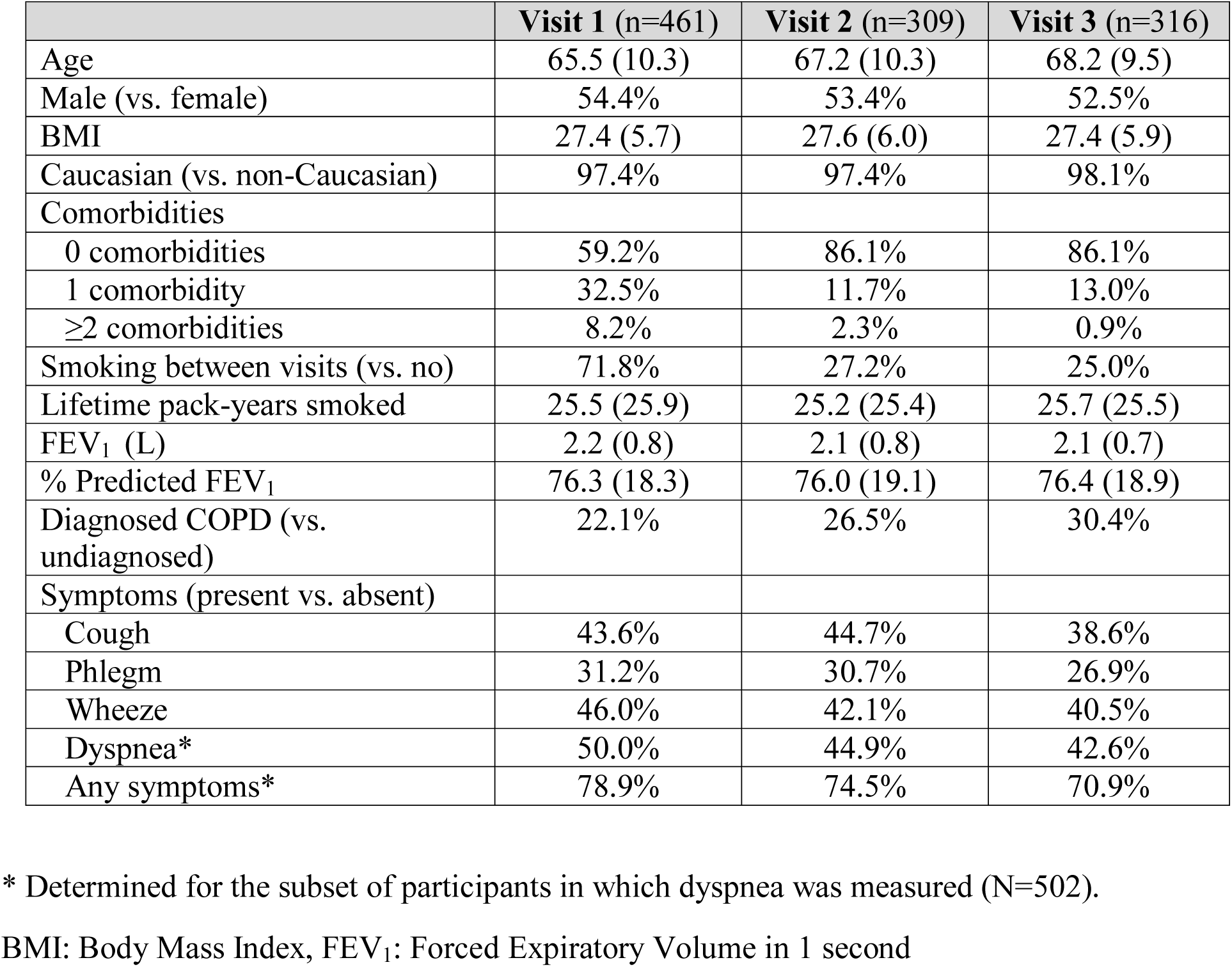
Characteristics of study participants at study visits. Means (and standard deviations) are reported unless otherwise indicated

### Objective 1: Heterogeneity in the occurrence of symptoms

Most participants did not report having cough, phlegm, wheeze, or dyspnea at each study visit, but only 18% of participants were completely asymptomatic throughout the study period. The asymptomatic participants tended to have mild airflow obstruction (mean of 89% predicted FEV_1_, 16% SD). The proportion of patients that reported a given symptom at least once during follow-up ranged from 38% for phlegm (the least common symptom) to 56% for dyspnea (the most common symptom). Symptoms were generally stable within participants: 66% of participants reported the same level of cough throughout their follow-up (the least stable symptom), and 75% for phlegm (the most stable symptom).

There was substantial variation in the individual-specific probabilities for the occurrence of symptoms (***Figure 1***). The median probabilities of an individual experiencing cough, wheeze, and dyspnea were 0.36, 0.34, and 0.44, respectively. In contrast, the median probability of experiencing phlegm was <0.01 and it was >0.99 for any symptoms. The interquartile range of probabilities was 0.13-0.68 for cough, <0.01-0.53 for phlegm, 0.10-0.72 for wheeze, 0.15-0.77 for dyspnea, and 0.56->0.99 for any symptoms. Median probabilities are depicted with blue lines and interquartile ranges are depicted with grey boxes in Figure 1.

**Figure 1.**
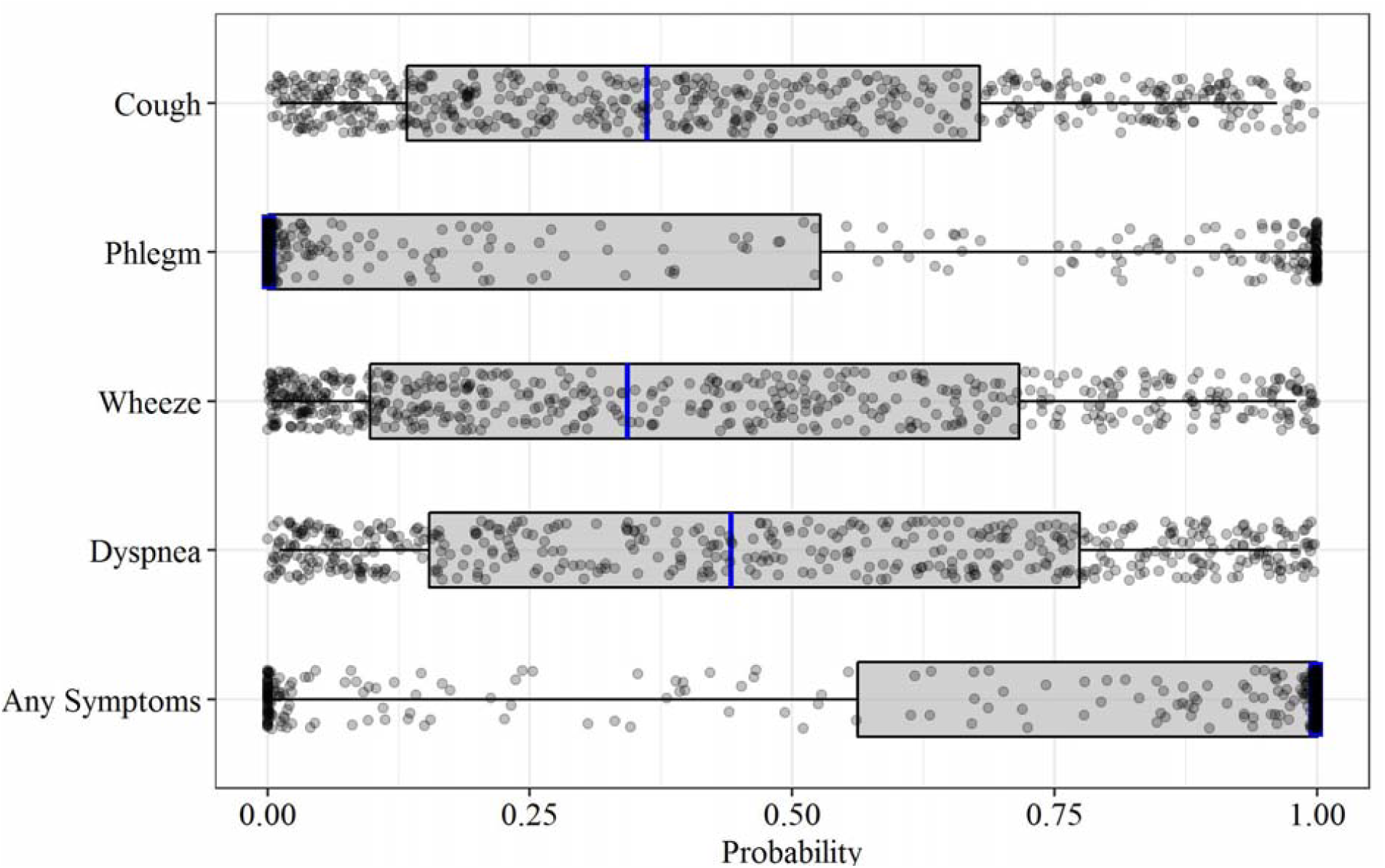
The distribution of individual-specific probabilities* of the occurrence of symptoms. The box spans the lower and upper quartiles (25%-75%) of individuals around the median (blue line). *Individual random effects are drawn from a normal distribution with a mean of 0 and standard deviation of the fitted random effects. The statistics shown by the boxes were determined from 1000 repetitions for each individual, and the points show the results of one repetition.

### Objective 2: Influence of lung function on symptom heterogeneity

The logistic regression models revealed relatively consistent associations between patient and disease characteristics and the presence of cough, phlegm, wheeze, dyspnea and any symptoms. Comparisons of the strength of associations across individual symptoms are shown in ***Figure 2***, and with any symptoms in ***Figure 3***. Lung function, sex, pack-years of smoking, BMI, and whether the participant had previously received a diagnosis of COPD were associated with most patient-reported symptoms. Ethnicity and the number of comorbidities were not associated with symptoms. Lung function was most strongly associated with the presence of any symptoms (OR per 100 mL increase in FEV_1_ 0.83, 95% CI 0.77-0.88), and least strongly associated with the presence of cough (OR 0.94, 95% CI 0.90-0.98). These results were similar when lung function was assessed as percent predicted FEV_1_ or GOLD grade in sensitivity analyses (results not shown). Higher pack-years of smoking, BMI, and a previous diagnosis of COPD were all associated with an increased odds of reporting most symptoms. Males were more likely than females to report the presence of cough (OR 2.12, 95% CI 1.18-3.81), phlegm (OR 6.95, 95% CI 2.74-17.59), wheeze (OR 2.86, 95% CI 1.34-6.11), and any symptoms (OR 3.16, 95% CI 1.38-7.23). Summer (vs. winter) was associated with increased reporting of cough (OR 2.02 95% CI 1.12-3.62), wheeze (OR 2.80 95% CI 1.44-5.46) and any symptoms (OR 2.89, 95% CI 1.34-6.24) when it was included in sensitivity analyses.

**Figure 2.**
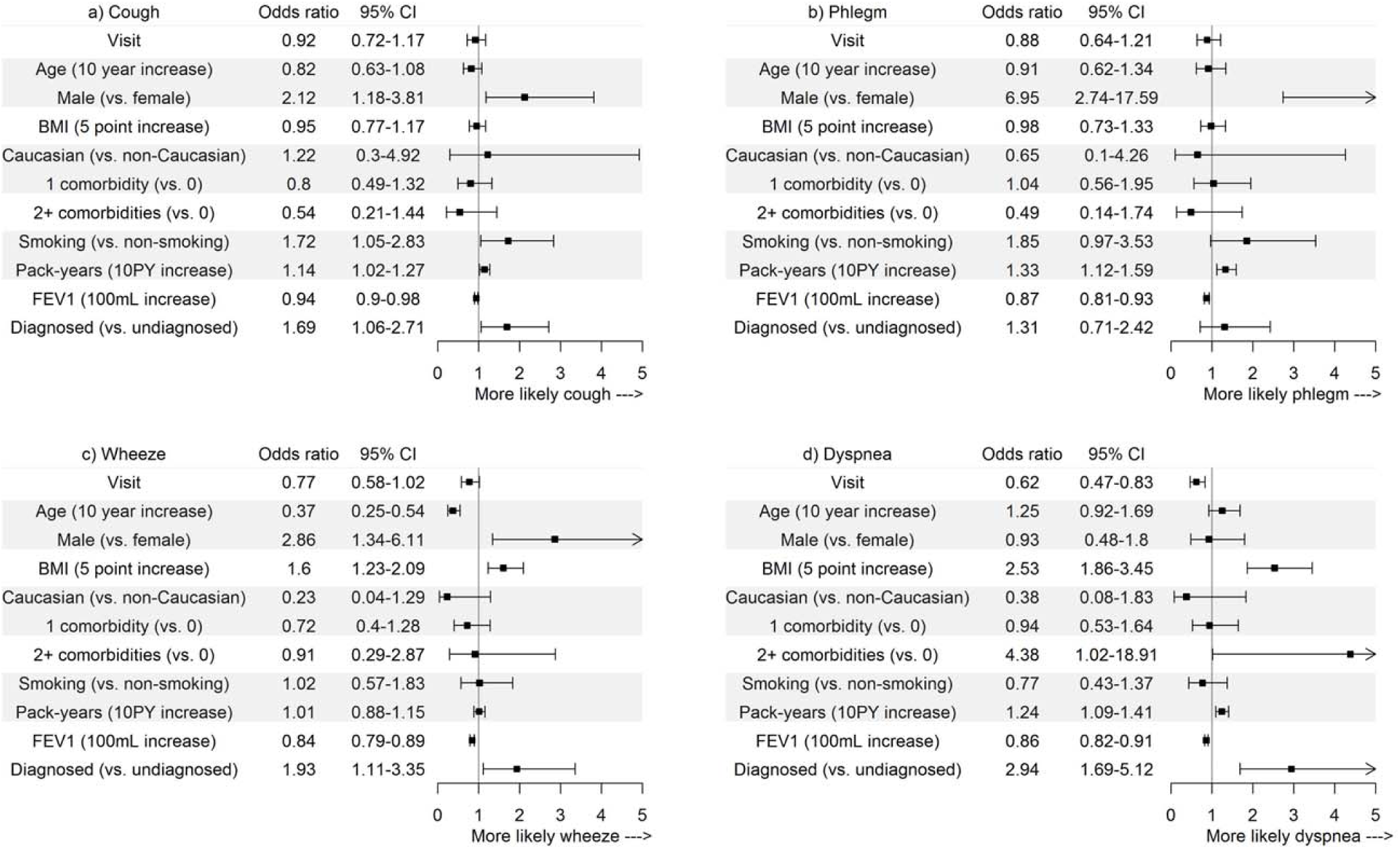
Odds ratios for the associations between independent variables and the presence of a) cough, b) phlegm, c) wheeze, and d) dyspnea.

**Figure 3.**
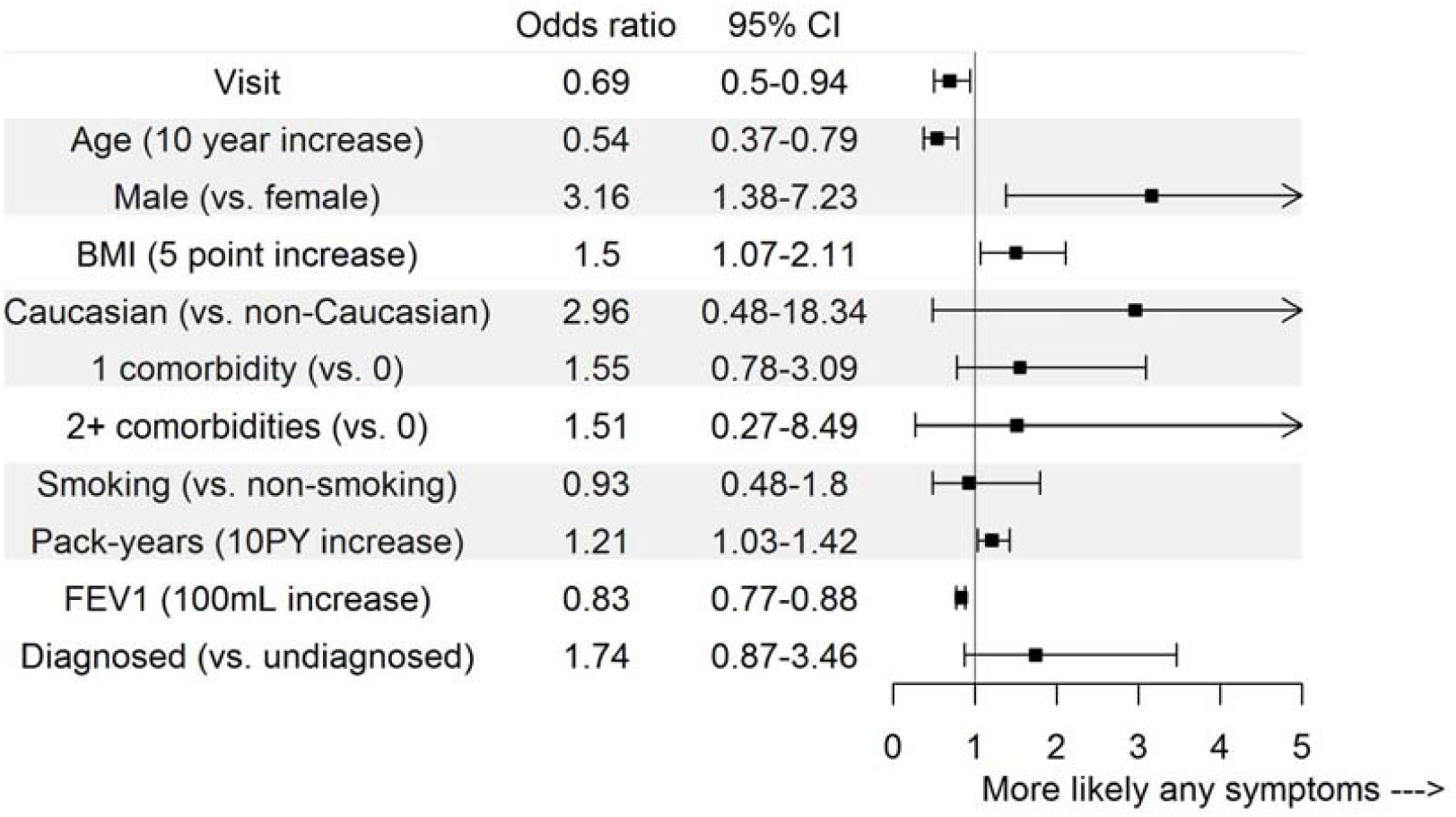
Odds ratios for the associations between independent variables and the presence of any symptoms.

The proportion of between-individual variation in the occurrence symptoms that could be attributed to participants’ measured characteristics (all independent variables in the full models) was 21%, 86%, <1%, 44%, and 92%, for cough, phlegm, wheeze, dyspnea, and any symptoms, respectively (***Table 2***). The proportion of variation explained by FEV_1_ alone ranged from 0% (for wheeze) to 82% (for phlegm, ***Table 2***).

**Table 2.** Percentage of between-individual variation in symptoms explained by individual’s lung function and all measured characteristics combined

|  | **Cough** | **Phlegm** | **Wheeze** | **Dyspnea** | **Any**<br>**Symptoms** |
| --- | --- | --- | --- | --- | --- |
| FEV<sub>1</sub> | 3% | 82% | 0% | 26% | 19% |
| All measured characteristics* | 21% | 86% | <1% | 44% | 92% |

\* Visit, Age, Sex, Caucasian, BMI, Comorbidities, Smoking status, Pack-years of smoking, Diagnosis status, FEV<sub>1</sub>
BMI: Body Mass Index, FEV<sub>1</sub>: Forced Expiratory Volume in 1 second

## 4.0 DISCUSSION

We have characterized heterogeneity in the occurrence of respiratory symptoms between patients with persistent airflow limitation and assessed the extent to which commonly measured patient and disease characteristics explained the observed heterogeneity in symptoms. Respiratory symptoms were very common in this sample of the general population; four out of every five participants reported experiencing symptoms despite over 90% of patients having mild to moderate COPD, and only 26% of them having been diagnosed with COPD. Dyspnea was the most common symptom, followed by cough and wheeze. Individual-specific probabilities for the occurrence of symptoms were highly variable between individuals and for different symptoms. The interquartile range of probabilities was the largest for dyspnea and wheeze, indicating greater variability between individuals in the presence of these symptoms than for cough and phlegm. For phlegm, the majority of individuals had a probability of experiencing phlegm near 0 or 1 (visible in Figure 1 as a higher density of points at the edges of the plot). In contrast, the individual-specific probabilities for cough, wheeze, and dyspnea were more evenly spread across the range of possible values. This indicates that phlegm is more stable in nature, and that individuals who do not currently have phlegm are unlikely to report it in the future. Indeed, a pan-European study reported that daily and weekly variability in dyspnea, wheeze, and cough were higher than that for phlegm.^25^ Our findings extend these observations on symptom variability within individuals to variability between individuals in the occurrence of symptoms. As a result, tools for assessing COPD severity that involve the measurement of symptoms (such as the GOLD ABCD assessment tool)^2^, are likely to be more or less variable over time, depending on the symptom measured.

The proportion of heterogeneity explained by the measured characteristics of participants differed substantially between symptoms. Most heterogeneity in the occurrence of phlegm and any symptoms, and approximately half the heterogeneity in dyspnea, was explained by the demographic and clinical characteristics of participants included in the models. In contrast, these characteristics explained very little heterogeneity in cough and wheeze, indicating that other characteristics not included in our models are more important drivers of these symptoms. Indeed, age, sex, BMI, smoking history, and lung function (cough only) were weakly correlated with cough frequency^26^ and the presence of wheezing^27^ in previous studies. Instead, cough frequency was driven by current smoking intensity and percentage of sputum neutrophils,^26^ and the presence of wheezing was associated with frequent exacerbations and increased dyspnea.^27^

Although lung function has traditionally been regarded as the primary driver of respiratory symptoms,^28^ we found that FEV_1_ explained the majority of between-individual variation in only the occurrence of phlegm, although it explained a substantial minority of variation in dyspnea and any symptoms. This finding is in line with the observation of high symptom variability within levels of disease severity,^29^ and high short-term variability in symptoms that is not due to changes in lung function.^4,12^ Our results extend these previous studies by examining the role of FEV_1_ in each symptom individually. They suggest that lung function is the primary driver of the occurrence of phlegm, an important but not dominant driver of the occurrence of dyspnea and any symptoms, but it explains very little between-individual variation in the occurrence of cough and wheeze.

In addition to analyzing heterogeneity in symptoms, we documented associations between symptoms and age, sex, BMI, smoking pack-years, FEV_1_, and a previous diagnosis of COPD. In particular, we observed substantial sex-based differences in the reporting of all symptoms apart from dyspnea. Controlled for disease severity, smoking history, and other variables, male patients were over three times more likely to be symptomatic, and seven times more likely to report experiencing phlegm. Whether this is a biological phenomenon, or due to gender-related differences in the experience of symptoms,^30^ remains to be further assessed.

Unique features of this study are its population-based sample and our assessment of the different sources of variation in symptoms. The associations determined from conventional regressions describe the relation between patient and disease characteristics and the presence of a symptom for an average participant. Our use of a random effect term in our models enabled us to extend these results by describing the extent to which these population-level associations apply to a given individual. We found that variation between individuals in the presence of phlegm and any symptoms was very well described by these population-level associations, but this was not the case for wheeze and cough (and somewhat the case for dyspnea). The assessment of variation at an individual-level is critical to fully characterizing heterogeneity in the natural history of COPD, and ultimately to enabling effective use of symptoms in risk prediction tools and case finding algorithms for COPD.

This study has several strengths. First, CanCOLD is a large, nationally representative sample of Canadians with COPD in the general population. The study employed standardized spirometry, validated questionnaires, and a long follow-up time. Our sample consisted primarily of patients with mild-moderate COPD, a population that is often underrepresented in large cohort studies. The population-based nature of the study makes it a better source for studying disease heterogeneity than clinical cohorts. However, this study also has several limitations. Patients reported their respiratory symptoms with a recall period that spanned the length of time between study visits, which could reach a maximum of three years. The long duration of the recall period is likely to have resulted in inaccuracies in symptom reporting. In addition, we only assessed the presence of symptoms, not their intensity. A more granular measurement of patient symptoms could provide a more nuanced assessment of symptom variability. Finally, we could not investigate the impact of factors that were not included in our model, in particular the use of treatment to control symptoms. Although this would likely have reduced the proportion of variation in symptoms that we attributed to unmeasured characteristics, the long recall period is less influenced by short-term variation in symptoms due to treatment.

## 5.0 CONCLUSION

We assessed a sample of the general population with mostly mild-moderate COPD and found substantial variation in the occurrence of respiratory symptoms between individuals. Lung function explained the majority of between-individual variation in only the occurrence of phlegm, and a much smaller proportion of variation in dyspnea, cough, and wheeze. For cough and wheeze in particular, commonly measured patient and disease characteristics explained very little heterogeneity in the occurrence of these symptoms. Overall, the observed differences in symptom variation may reflect the divergent etiology of symptoms associated with COPD, and suggests that defining endotypes that are predictive of a high symptom burden is a key area of future research.

## ABBREVIATIONS

COPD: Chronic Obstructive Pulmonary Disease
CanCOLD: Canadian Cohort of Obstructive Lung Disease Study
FEV_1_: Forced Expiratory Volume in 1 second
FVC: Forced Vital Capacity
GOLD: Global Initiative for chronic Obstructive Lung Disease
LLN: Lower Limit of Normal
MRC: Medical Research Council dyspnea scale
OR: Odds Ratio
SD: Standard Deviation

## DECLARATIONS

### Ethics approval and informed consent

Ethics approval for CanCOLD was obtained from the relevant institutional review board at each study site. Written informed consent was obtained from all participants prior to study entry.

### Consent for publication

Not applicable.

### Availability of data and materials

The data analyzed in the current study are not publicly available but may be made available from the CanCOLD Research Group upon reasonable request.

### Author Contributions

WT and JB are co-Principal Investigators of the CanCOLD study. MS and KJ formulated the current study idea. KJ performed all data analyses and wrote the first draft of the manuscript. AS provided guidance on the statistical analysis. All authors contributed to interpretation of findings, critically commented on the manuscript and approved the final version. MS is the guarantor of the manuscript.

## Acknowledgements

The authors thank the men and women who participated in the study and individuals in the CanCOLD Collaborative Research Group.

Members of the CanCOLD Collaborative Research Group are as follows. Executive Committee: Jean Bourbeau (McGill University, Montreal, Canada); Wan C. Tan, J. Mark FitzGerald; Don Sin (UBC, Vancouver, Canada); Darcy Marciniuk (University of Saskatoon, Saskatoon, Canada); Dennis E. O’Donnell (Queen’s University, Kingston, Canada); Paul Hernandez (Dalhousie University, Halifax, Canada); Kenneth R. Chapman (University of Toronto, Toronto, Canada); Robert Cowie (University of Calgary, Calgary, Canada); Shawn Aaron (University of Ottawa, Ottawa, Canada); F. Maltais (University of Laval, Quebec City, Canada). International Advisory Board: Jonathon Samet (Keck School of Medicine of USC, Los Angeles, CA); Milo Puhan (John Hopkins School of Public Health, Baltimore, MD); Qutayba Hamid (McGill University, Montreal, Canada); James C. Hogg (UBC James Hogg Research Center, Vancouver, Canada). Operations Center: Jean Bourbeau (PI), Carole Jabet, Palmina Mancino, (McGill University, Montreal, Canada); Wan C. Tan (co-PI), Don Sin, Sheena Tam, Jeremy Road, Joe Comeau, Adrian Png, Harvey Coxson, Jonathon Leipsic, Cameron Hague (University of British Columbia James Hogg Research Center, Vancouver, Canada). Economic Core: Mohsen Sadatsafavi (University of British Columbia, Vancouver, Canada). Public Health Core: Teresa To, Andrea Gershon (University of Toronto, Toronto, Canada). Data Management and Quality Control: Wan C. Tan, Harvey Coxson (UBC, Vancouver, Canada); Jean Bourbeau, Pei Zhi Li, Zhi Song, Yvan Fortier, Andrea Benedetti, Dennis Jensen (McGill University, Montreal, Canada). Field Centers: Wan C. Tan (Vancouver PI), Christine Lo, Sarah Cheng, Elena Un, Cindy Fung, Nancy Haynes, Junior Chuang, Licong Li, Selva Bayat, Amanda Wong, Zoe Alavi, Catherine Peng, Bin Zhao, Nathalie Scott-Hsiung, Tasha Nadirshaw (UBC James Hogg Research Center, Vancouver, Canada); Jean Bourbeau (Montreal PI), Palmina Mancino, David Latreille, Jacinthe Baril, Laura Labonté (McGill University, Montreal, Canada); Kenneth Chapman (Toronto PI), Patricia McClean, Nadeen Audisho (University of Toronto, Toronto, Canada); R. Cowie and B. Walter (Calgary PI), Ann Cowie, Curtis Dumonceaux, Lisette Machado (University of Calgary, Calgary, Canada); Paul Hernandez (Halifax PI), Scott Fulton, Kristen Osterling (Dalhousie University, Halifax, Canada); Shawn Aaron (Ottawa PI), Kathy Vandemheen, Gay Pratt, Amanda Bergeron (University of Ottawa, Ottawa, Canada); Denis O’Donnell (Kingston PI), Matthew McNeil, Kate Whelan (Queen’s University, Kingston, Canada); François Maltais (Quebec PI), Cynthia Brouillard (Université Laval, Quebec City, Canada); Darcy Marciniuk (Saskatoon PI), Ron Clemens, Janet Baran (University of Saskatoon, Saskatoon, Canada).

